# Colour vision and information theory: the receptor noise-limited model implies optimal colour discrimination by opponent channels

**DOI:** 10.1101/2020.08.07.242065

**Authors:** Sebastián Risau-Gusman

## Abstract

In order to interpret animal behaviour we need to understand how they see the world. As colour discrimination is almost impossible to test directly in animals, it is important to develop theoretical models based in the properties of visual systems. One of the most successful is the receptor noise-limited (RNL) model, which depends only on the level of noise in photoreceptors and opponent mechanisms. Here optimal colour discrimination properties are obtained using information theoretical tools, for the early stages of visual systems with and without colour opponent mechanisms. For most biologically relevant conditions the optimal discrimination function of an ideal observer coincides with the one obtained with the RNL model. Many variants of the model can be cast into the same framework, which permits meaningful comparisons across species. For example, it is shown that the presence of opponency seems to be the preferred hypothesis for bees, but not for budgerigars. Since this is a consequence of the presence of oil droplets, this could also be true for most other species of birds.

## 1 Introduction

For many animals colour vision is essential for activities as diverse as foraging and mate selection. The diversity of existent visual systems makes it very difficult to infer their perception of colour from what we know from humans. Furthermore, even when behavioural tests can be conducted, practical considerations impose a stringent limit on the number of stimuli that can be used. A complementary approach is to use our knowledge about vision in animals to infer colour discrimination properties, by building a theoretical model.

Visual processing starts with the absorption of photons by pigments inside photoreceptors, from where information is transferred across many stages to higher level processing neurons in the brain (in the visual cortex in mammals, or in the protocerebrum in some insects [1]). Given the complexity of this processing, a detailed mathematical model of colour discrimination is infeasible, as it requires the specification of parameters whose values are unknown. A “good” model should provide a reasonable good match for behavioural observations using only a few features of the visual system. In this sense, one of the most successful models of colour discrimination is the receptor noise limited (RNL) model [2, 3].

In this model, only the early stages of visual processing are considered and noise is only present at the level of photoreceptors and not in the opponent mechanisms. The achromatic part of the signal is also disregarded. The model uses distances defined in opponent mechanism space and thresholds for discrimination are derived from them. Both the metric of the space and the discrimination thresholds are given arbitrary definitions, but they seem reasonable and, most importantly, lead to a simple and intuitive model. It has been found to be very effective to predict spectral sensitivities of many different animals, such as honeybees [2, 3], budgerigars [4, 5] and triggerfishes [6].

A different approach to model construction is to assume that, given the constraints of its visual system, the observer can perform optimal discrimination or detection. In a more general framework this is known as *ideal observer theory* [7]. In order to analyze separately the contributions of the different stages of visual processing, it is useful to think that each stage processes information from downstream stages and provides information that has to be decoded by the processing centers upstream. One is then interested in undestanding the consequences of optimal upstream decoding for color discrimination. To assess optimality one can use maximum likelihood estimators [8, 9] or the Cramér-Rao bound [10] which gives a bound for the performance of all estimators. The discrimination function obtained is necessarily a lower bound for the real one, since further processing cannot improve discrimination because no new information is available. However, given that some parameters of the model are hard to obtain, one is more interested in obtaining a good approximation for the shape of the discrimination function. The function obtained using only the number of photons absorbed has been shown to provide a reasonable approximation for the real one in human subjects [8, 9, 10].

Here this approach is taken one step further by considering the effect of opponent mechanisms, as well as the effect of noise and chromatic adaptation in the output of the photoreceptors. When the intensity of the stimuli is not too small, the discrimination function is shown to reduce to a simple form, that coincides with the function given by the RNL model. The common framework of Information Theory allows us to study other variants, whose predictions can then be meaningfully compared with those of the RNL model.

## 2 Model of photon absorption and photoreceptor output

Visual systems have different types of photoreceptors, with spectral sensitivities *R*_*i*_(*λ*), where *i* is the type of photoreceptor. These sensitivities should take into account what happens before light reaches the photopigments (i.e. absorptions by ocular media, screening, etc.). For simplicity, in most of the following three types of photoreceptors will be used for the calculations, but the results are easily extended to more types. Trichromat photoreceptors are usually designed as *S, M* and *L*, with sensitivities peaking at wavelengths *λ*_*S*_ *< λ*_*M*_ *< λ*_*L*_.

The number of photons, *k*, of wavelength *λ*, emitted in a given time by most of the commonly used light sources with intensity *I*(*λ*) follows a Poisson distribution with average *I*(*λ*). The number of these photons absorbed by a cone depends on both its spectral sensitivity and its cross sectional area *β*_*i*_. The probability that a cone of type *i* (*i* = *S, M, L*) absorbs *k*_*i*_ photons of a monochromatic source is a Poisson distribution of average *R*_*i*_(*λ*)*β*_*i*_*I*, where *I* is now the intensity of the source per unit surface at the position of the cone. For a general light source, this probability must be integrated over all possible wavelengths, and the result is [10]:

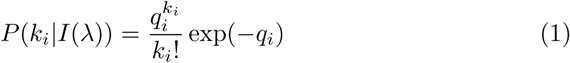

Where

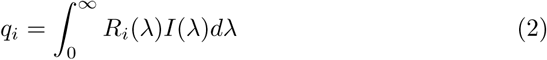

*q*_*i*_ is usually called the *quantum catch* of photoreceptor *i*. Note however that this is not the actual number of photons absorbed by the cone, which is a random number, but its average.

The absorption of a photon starts the process of phototransduction whereby an electrochemical signal is generated. The basic mechanisms are highly conserved both in vertebrates and invertebrates [11], and lead to a significant amplification of the light signal [12]. Even though amplification is not as large as in rods [13], it is still large in cones, and may be considered one of the most important sources of noise.

If no light adaptation is present, *x*_*i*_, the output of cone *i*, can be modelled as the product of the number of photons absorbed and a noisy amplification term: *x*_*i*_ = *k*_*i*_*αψ*_*i*_, where *ψ*_*i*_ is drawn from a Gaussian distribution with mean 1 and variance *σ*_*i*_, *α* is the mean amplification, and *k*_*i*_ has the distribution given in Eq. 1. As most visual systems process stimuli with intensities spanning at least 7 orders of magnitude, it is reasonable to expect some degree of light adaptation in photoreceptors. This is the case for many animals (e.g. primates [14], turtles [15] and fruit flies [16]): the cone photocurrent saturates when light intensity is either too high or too low, and in the middle region it has an approximately logarithmic dependence with intensity. In this intermediate regime, known as Weber-Fechner law [17], the output of the photoreceptor mechanisms can be written as:

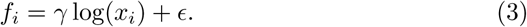

The value of constants *α, γ* and *ϵ* is left unspecified, because they are not necessary for the determination of the threshold function, as shown below.

## 3 Colour discrimination and sensitivity

In humans, wavelength discrimination is usually tested by using two identically illuminated fields, and asking the observer to change the wavelength of one of them until that field is perceived as having a different colour from the reference. In this case, both stimuli have the same intensity and the relevant magnitude is the wavelength *λ*. In animals, colour vision has to be tested indirectly, and there are different techniques for this [18]. In detection experiments, they consist of two steps. In the first, the animal is trained to associate a reward with an object which has been painted or illuminated with the colour to be tested. In the second step, the rewarded object, illuminated with different intensities, is presented with with a similar, unilluminated reference object and the choice made by the animal is registered. The relevant information is *P*_*error*_, the fraction of times that the rewarded object was not chosen. As the difference between the two objects is increased, this fraction moves from 0.5 (objects cannot be distinguished) to 0 (objects can be perfectly distinguished). Usually, the discrimination threshold is set at 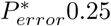.

In the discrimination experiments mentioned above, two different stimuli are presented and the observer has to decide which is the rewarded one. This is known as a *two-alternative forced choice* task in the psychophysics literature [19]. The quantities (such as intensity or wavelength) that define these stimuli are noted as *s* and *s* + *δs*, and are unknown to the observer, whose task is to decide which stimulus is the rewarded object. Performance is characterized by the fraction of errors, which is a monotonically decreasing function of *δs*. The processing of the two stimuli generates stochastic *responses* r_1_ and r_2_, which the inner *decoder* must use to perform the discrimination task. r can be the number of photons absorbed by the cones, the corresponding photocurrents, or the output of the opponent mechanisms. When noise is present both in the stimulus and the detection system, the responses are not the same in different trials of an experiment. The probability of r when a stimulus *s* has been presented is *P* (r|s), usually called *likelihood* of *s*. One of the methods that the decoder can use for discrimination is the Maximum Likelihood method [20], which consists in choosing the stimulus whose response gives the largest value of *P* (r|s + *δ*s) (assuming that the rewarded object has magnitude *s*+*δs*). If the responses have Gaussian distributions, or there are many uncorrelated units, the probability of a discrimination error is

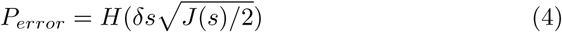

where 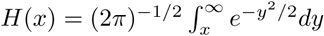, and *J*(*s*) is the Fisher information:

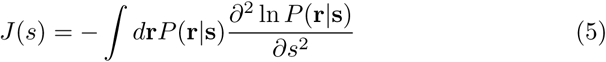

This is optimal when the number of response units is large. Another method is based on the existence of an internal *estimator*. At each trial, the estimator gives estimates *ŝ*_1_ and *ŝ*_2_ of the magnitudes of the stimuli presented. The stimulus chosen as the rewarded is the one with the largest value of *ŝ*. Because of stochasticity, the values of *ŝ* are not the same in each trial, but have probability distributions *g*_*i*_(*ŝ*) (*i* = 1, 2) that are unknown to the experimenter. It is usually assumed that estimates are *unbiased*, i.e. that their mean values are equal to the respective stimuli. Evidently, the performance of the classifier will depend on how concentrated the *g*_*i*_(*ŝ*) are about about their mean value. If *δ*_*s*_ is small, it can be assumed that *g*_1_(*ŝ*) = *g*_2_(*ŝ*) = *g*(*ŝ*). Furthermore, if *g*(*ŝ*) is a Gaussian distribution with a variance *σ*_2_, it can be shown [21] that, for the two-alternative forced choice task, 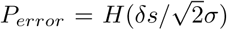. The Cramér-Rao bound states that, in general, the variance of *g*(*ŝ*) must satisfy *σ*_2_ ≥ 1*/J*(*s*). For the optimal estimator the equality holds, and if it is further assumed that *g*(*ŝ*) is Gaussian one recovers Eq 4. In order to define a discriminability or detection threshold, a value 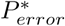 must be chosen, such that the stimulus is considered to have been detected or discriminated when it was not chosen in at most 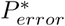 of the trials. For an observer implementing optimal estimation (i.e an ideal observer) the threshold stimulus is:

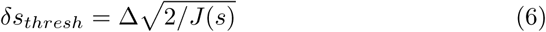

where Δ is the solution of the equation:

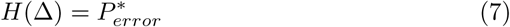

For 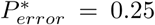, we have Δ = Δ_*G*_ ≈ 0.67. Even though *g*(*ŝ*) might not be Gaussian, for visual systems it seems reasonable to assume that the animal makes the decision averaging the estimations made during several integration times (≈ 10 − 20 ms.). The distribution of such averages would be very close to a Gaussian, because of the Central Limit Theorem. Because of the intrinsically noisy nature of the processing steps upstream, there will be some loss of information, resulting in discrimination thresholds larger than predicted by Eq. 6. It will be further assumed that the amount of information lost is independent of *s*. Thus, the hypothesis is that the dscrimination thresholds observed should be given by Eq. 6, but the parameter Δ must be obtained by fitting discrimination data for each animal, and Δ_*G*_ should be considered as a lower bound to the possible values of Δ.

In the case of monochromatic stimuli characterized by wavelength and intensity, two different types of tests can be performed. In colour discrimination tests, the objects are slightly different in color, or are illuminated with lights of slightly different wavelengths, whereas in detection tests the difference lies in the intensity of the illumination. In the following it will be assumed that stimuli are of the form

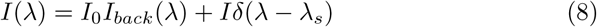

where *I*_*back*_(*λ*) is the normalized spectrum of the background, and *δ*(*λ*) (a very narrow function concentrated at *λ*_*s*_) is the normalized spectrum of the stimulus, with *I*_0_ and *I* their respective total intensities. Using this, the average number of photons absorbed by a photoreceptor of type *i*, usually called the coordinate *i* of the stimulus in receptor space, is:

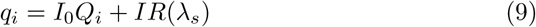

where *Q*_*i*_ are the receptor coordinates of the background.

## 4 Fisher information for early visual processing

In order to obtain the discrimination properties that can be inferred from the output of the photoreceptors, it will be assumed that r = (*f*_*S*_, *f*_*M*_, *f*_*L*_), with the *f*_*i*_ defined in Eq. 3. One important property of the Fisher information is that it does not change if the responses are replaced by monotonous functions of them. Thus, *P* ((*f*_*s*_, *f*_*M*_, *f*_*L*_)|*s*) can be replaced by *P* ((*x*_*s*_, *x*_*M*_, *x*_*L*_)|*s*). Furthermore, in order to obtain an analytical expression, the probability distribution of *x*_*i*_ is approximated by a Gaussian distribution with mean value *q*_*i*_ and variance 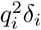 with 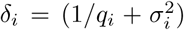. For photopic vision (*q*_*i*_ ≳ 100 photons) and low photoreceptor noise, this distribution is very close to the true distribution of *x*_*i*_ (see Fig.1 in the Supplementary Material). In fact, the difference between the distributions is proportional to *δ*_*i*_ (Section A in the Supplementary Material). Within this model, the signal to noise ratio (SNR) for an isolated photoreceptor in bright light is 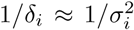. Thus, it depends mainly on the amplification process and 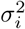 can be well approximated by the reciprocal of SNR that is measured in isolated photoreceptors in bright light.

Using *P* ((*x*_*s*_, *x*_*M*_, *x*_*L*_) | *s*) in Eq. 5, the Fisher information provided by the photoreceptor output about the monochromatic stimulus is

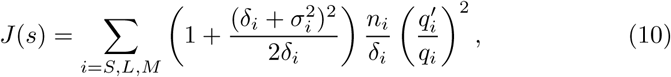

where *s* can be either intensity or wavelength of the stimulus, *n*_*i*_ is the number of photoreceptors of type *i* in the receptive field, and 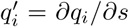. Given that for photopic illumination the SNR is in general relatively large [3, 22], it will be assumed that *σ*_*i*_ ≪ 1. In this regime the term within parentheses is close to 1, and the Fisher information can be well approximated by

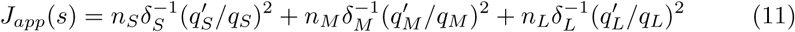

When intensity thresholds are needed, *J*(*λ*) is calculated using 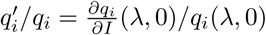 whereas for discrimination wavelengths, *J*(*λ*) is calculated using 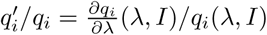.

## 5 Fisher information for opponency channels

For *n* photoreceptor types, there are *n*(*n* − 1)*/*2 possible channels that oppose all possible pairs of photoreceptors. The output of the *ij* channel is modelled as:

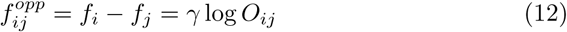

with *O*_*ij*_ = *x*_*i*_*/x*_*j*_. Note that the outputs of all the possible channels are not independent. For example, the output of channel 13 can be obtained from 12 and 23 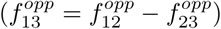. The largest set of independent channels can have at most *n* − 1 channels.

As mentioned, the Fisher information is the same for any monotonous function of the responses. Thus, in order to simplify calculations, the variables *O*_*ij*_ are used. Using the Gaussian approximation of the previous section, the probability distribution for *O*_*SM*_ and *O*_*LM*_ can be calculated exactly (see Appendix). The calculation of *J*(*s*) is straightforward but the expression obtained is long and contains integrals that can only be evaluated numerically (see Section B of the SM). However, for photopic conditions and relatively sma<u>l</u>l receptor noise, the expression becomes much simpler: 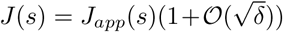, where 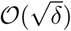 represents terms that vanish as 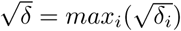 (Section B of the SM) and

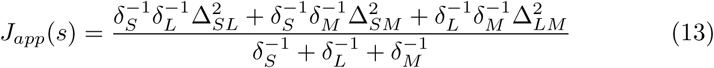

With

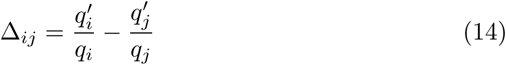

with 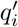 as in the previous section (*i, j* = *S, L, M*). For realistic values of photoreceptor noise and photopic illumination, the approximation is virtually undistinguishable from the exact expression of *J*(*s*) (see Fig. 1). Combining Eqs. (13), (6), and (9), the discrimination function obtained is exactly the same as in the RNL model when each opponency channel compares the signal of two single photoreceptors [3].

**Figure 1:**
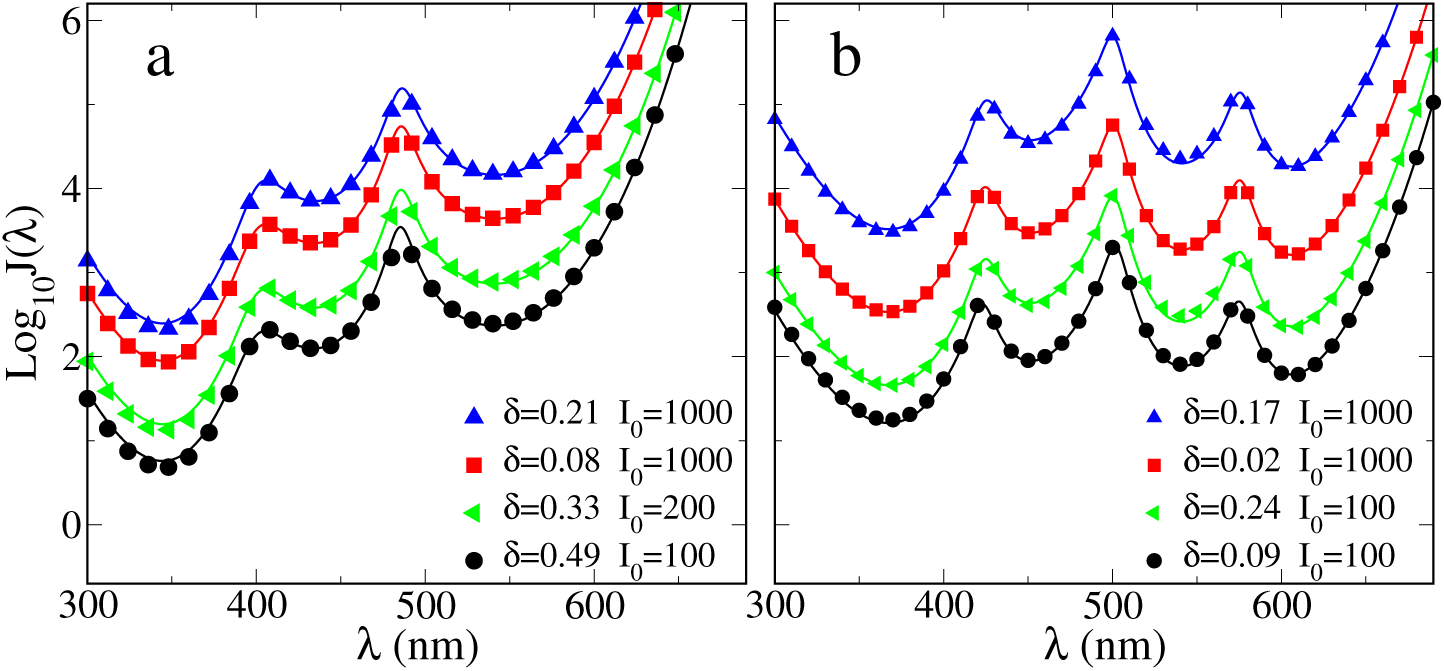
Comparison of exact Fisher information and its approximation for several levels of noise, in two different opponent systems. a) Trichromat (using data from honeybees), b) Tetrachromat (using data from budgerigars). Symbols represent Fisher information, whereas lines represent the corresponding approximations

In the more general case, a receptive field contains *n*_*S*_, *n*_*M*_, and *n*_*L*_ photoreceptors of types *S, M* and *L* respectively, and the opponent channels may weigh differently positive and negative contributions. The output of the opponent channel *ij* is in this case,

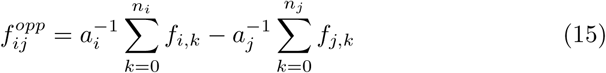

where *f*_*i,k*_ is the output of the *k*-th receptor of type *i*, and *a*_*i*_ is the weight assigned to the input from the *n*_*i*_ photoreceptors of type *i*. When *a*_*i*_ = *n*_*i*_ the channels are called *purely chromatic* because the output is completely independent from the achromatic part of the stimulus. In other words, channel output does not depend on the *total* intensity of the illumination.

In order to calculate the Fisher information it is useful to define the new variables 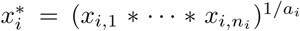. They allow us to write the output of the channel as 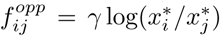. In order to get an analytical expression for the Fisher information, the distribution of the random variables 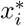 is again approximated by a Gaussian, now with mean 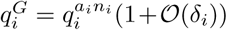 and variance 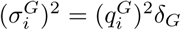 with *δ*_*G*_ = (*δ*_*i*_*/n*)(1+𝒪(*δ*_*i*_)). Again, the difference between the exact distributions and the approximations tends to 0 in the limit of vanishing *δ*_*i*_. Furthermore, the approximation is most effective in the case of purely chromatic channels (see Section C of the SM). Using these approximations, the dominant term of *J*(*s*) can be obtained from Eq. 13, replacing *q*_*i*_ by 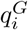 and *δ*_*i*_ by 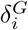. The resulting expression is:

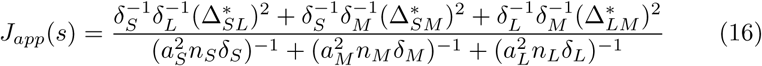

Where

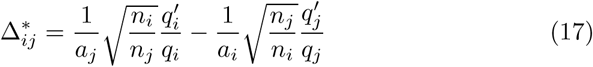

When *a*_*i*_ = *n*_*i*_ this is exactly the expression given by the RNL mode for a general receptive field [2, 3]. Eq. 16 applies to any opponency channels that can be written as a linear combination of binary opposition channels. Consider, for example, the channels S-(L+M) and L-M, that seem to be common in mammals. The sensitivity of these channels is the same as for channels L-M and S/2-M. In other words, the discrimination properties are obtained from Eq. 16 using *a*_*S*_ = 1*/*2, *a*_*M*_ = 1 and *a*_*L*_ = 1. In particular, any set of purely chromatic opponent channels can be written as a combination of binary opponent channels.

## 6 Comparison of systems with and without opponency

The Fisher information approach provides a framework within which systems with and without opponency can be meaningfully compared. It has been shown that the RNL model gives a good description of the colour vision of honeybees (*Apis mellifera*) [2, 3] and budgerigars (*Melopsittacus undulatus*) [4, 5]. Fig. 2 shows the sensitivities measured for three different bees in two sets of experiments [23, 3]. Three different sensitivity functions have been fitted to the data: without opponency channels, with purely chromatic opponency channels, and with non purely chromatic opponency channels (using *a*_*i*_ = 1 for *i* = *S, L, M*). The results of the fit are given in Table 1. The curves and the table show that purely chromatic channels (i.e. the RNL model) describe the data much better than the other mechanisms. For the bee in Fig. 2d there is some discrepancy for long wavelengths, but in [23] it is explicitly mentioned that the results for these wavelengths are somehow anomalous. But even in this case the RNL model is the only one which displays a large dip at a frequency close to 500 nm., a dip that has been observed for most bees in both works [23, 3].

**Table 1:** Values of Δ and *R*_2_ obtained for the best fits of three different sensitivity functions to the data of three bees. The functions tested correspond to systems with no opponency (NoOpp), with purely chromatic opponency channels (PC), and with non purely chromatic opponency channels (NPC).

|  | Bee 10 [3] |  |  | Bee 25 [23] |  |  | Bee 15 [23] |  |  |
| --- | --- | --- | --- | --- | --- | --- | --- | --- | --- |
|  | NoOpp | PC | NPC | NoOpp | PC | NPC | NoOpp | PC | NPC |
| $\Delta$ | 2.47 | 1.44 | 1.39 | 1.18 | 1.00 | 0.92 | 3.15 | 2.88 | 2.97 |
| $R^2$ | 0.926 | 0.992 | 0.838 | 0.974 | 0.984 | 0.848 | 0.953 | 0.948 | 0.724 |

**Figure 2:**
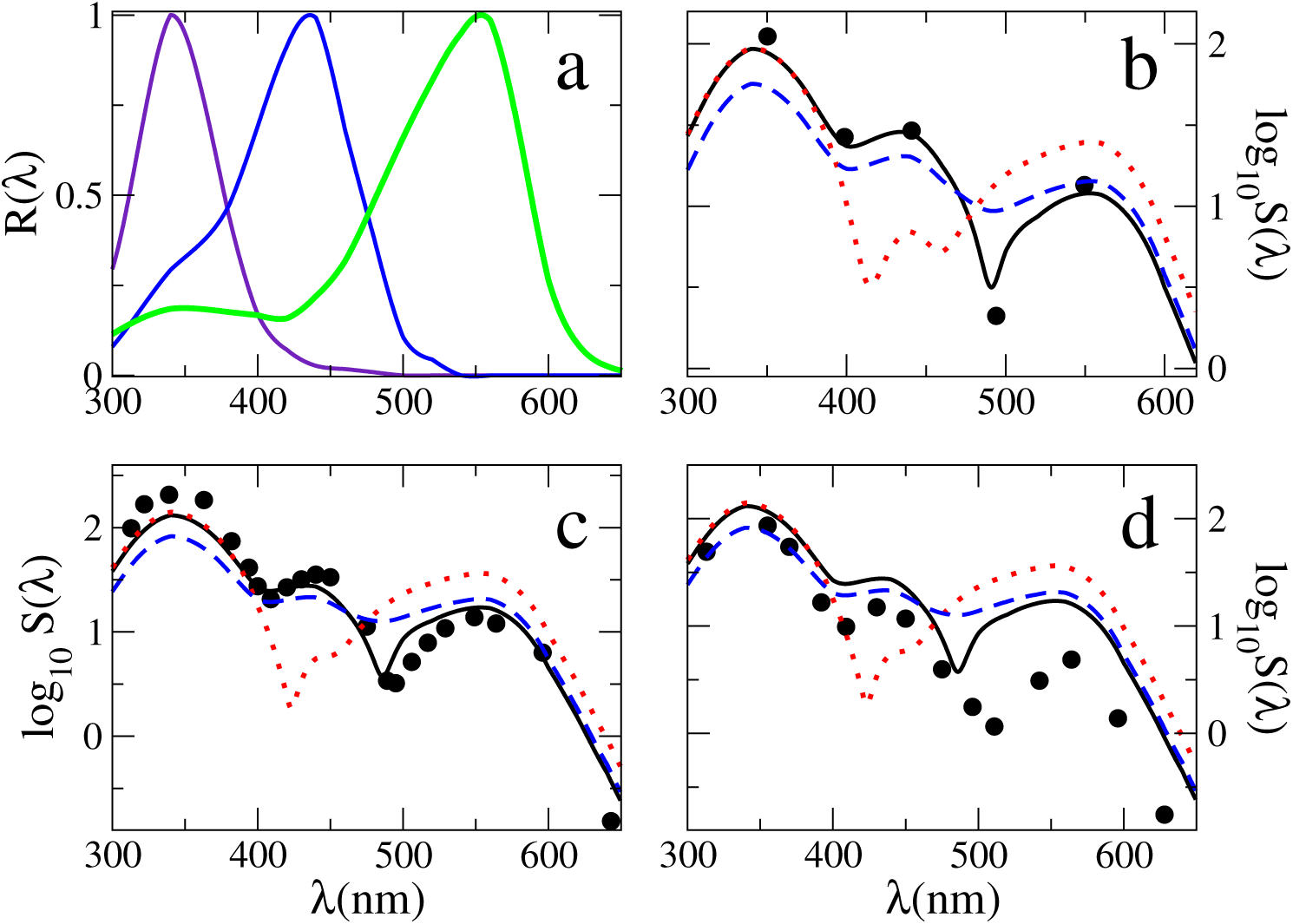
Best fits of sensitivity functions to data from three different bees. a) Normalized absorbance spectra of bee photoreceptors (S: violet, M: blue, L: green curve). b), c) and d): Symbols give the relative sensitivity for three different bees, b) bee 10 [3], c) bee 25 [23], d) bee 15 [23]. Curves are theoretical sensitivities calculated for systems with: no opponency (dashed), purely chromatic opponency channels (full), non purely chromatic opponency channels (points).

On the other hand, for budgerigars the situation seems to be different. In [4, 5] it has been shown that the RNL model gives a good fit for the sensitivity of these parrots. However, when one compares the sensitivity functions for ideal observers with and without opponency channels, they turn out to be very similar (see Fig. 3). Even though they are not identical, the dispersion of the data obtained in experiments with budgerigars makes it almost impossible to decide which is the best model for the data. Of course, this does not rule out the existence of opponency in budgerigars. However, the argument of parsimony should tip the scale in favour of a system without opponency channels.

**Figure 3:**
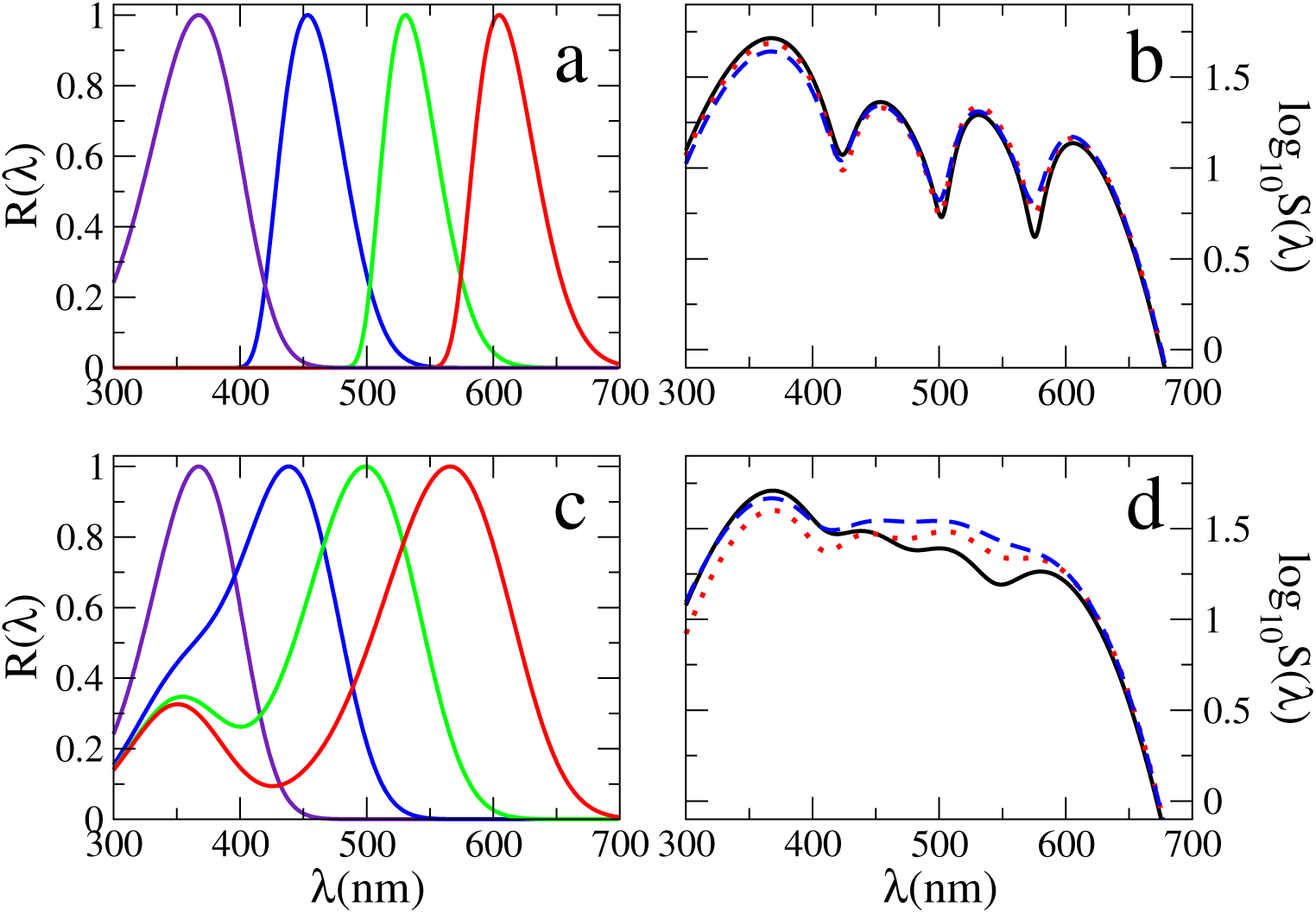
Comparison between models based on budgerigar photoreceptors. a) and c) Normalized absorbance spectra of budgerigar photoreceptors (UV: violet, S: blue, M: green, L: red) with (a) and without (c) oil droplets. b) and d): Theoretical sensitivities for systems with: no opponency (dashed), purely chromatic opponency channels (full), non purely chromatic opponency channels (points), and photoreceptors as in a) and c), respectively.

In a way, this was to be expected [24] because the effective absorbance spectra of the four photoreceptors have very small overlap (see Fig. 3a). Interestingly, this happens because the photoreceptors are screened by oil droplets.

Without these droplets the absorbance spectra would have a much larger overlap, and the differences between the sensitivity curves for the three mechanisms would be large enough to provide significantly different fits to the experimental data (see Fig. 3d).

## 7 Conclusion

The RNL Model [2] has been widely acknowledged to provide a good description of colour discrimination across many species. Originally, it was derived using a heuristic approach where a colour space and a metric inside it are chosen, and distances of colour loci are related to perception. Even though the agreement with the data shows that the choices made were very sensible, they were ultimately arbitrary. Other choices were possible, but the resulting models were nowhere as successful (see e.g. [25, 26, 27]). In the sections above colour discrimination properties have been derived using a completely different approach, based on Information theory. Interestingly, even though the resulting discrimination function is complicated, in the limit of bright illumination and low photoreceptor noise it can be written as the RNL discrimination function (with shot noise added) with correction terms that are negligible for all practical purposes.

The Fisher information approach allows a meaningful comparison of the RNL model with other variants corresponding to different visual systems. The interpretation of this comparison becomes straightforward, since the assumptions and approximations used to derive each of the models are exactly the same. Such comparisons are often needed to supplement goodness of fit arguments. For example, the fact that the RNL model gives a good fit for colour discrimination data of bees and budgerigars does not guarantee that the existence of opponency channels is the best hypothesis. Opponency seems indeed to be the preferred hypothesis for bees, but for budgerigars the absence of opponency is a hypothesis at least as strong as its presence. As this is a consequence of the presence of oil droplets, a widespread feature in birds [28], the lack of opponent channels might well be the preferred hypothesis for most avian species.

The present derivation provides a convenient framework to propose and analyse extensions of the RNL model. One possibility is to study the effect of adding noise to opponent channels, but there is almost no information about this feature in animals. A more interesting possibility is to study colour discrimination in low levels of illumination. It is important to note that in this limit at least one of the approximations used in this paper breaks down. However, the colour discrimination function could be calculated by dropping some of the approximations and resorting to numerical methods and the results can be compared with experimental studies where colour sensitivity is studied for different levels of background illumination [5].

## Acknowledgments

This work was partly supported by the ANPCyT, Argentina [grant number PICT 1042-2016].

