## Supplemental Material for "Colour vision and information theory: the receptor noise-limited model implies optimal colour discrimination by opponent channels"

### Supplementary Material for Color coding and information theory: the receptor noise-limited model implies optimal color discrimination by opponent channels


Sebastián Risau-Gusman *

Consejo Nacional de Investigaciones Científicas y Técnicas,
Centro Atómico Bariloche,
San Carlos de Bariloche, 8400 Río Negro, Argentina



**Abstract**

Here are shown the calculation of the Fisher information, and a sketch of the proofs the convergence of the approximations used in the paper.


#### A Approximation for photoreceptor outputs without adaptation

Let $x = k\psi$, where $\psi$ is drawn from a Gaussian distribution with mean 1 and variance $\sigma$, and $k$ is drawn from a Poisson distribution of mean $q$. $x$ has mean $q$ and variance $\sigma^2(q^2 + q) + q = \sigma^2\delta$, with $\delta = \sigma^2 + 1/q + \sigma^2/q \approx \sigma^2 + 1/q$, as in the main text. $\delta$ has been approximated because we are interested here in photopic illumination, where $q \ll 1$.

Since the exact distribution of $x$ can only be calculated numerically, we wish to approximate it by a distribution that can be written in closed form. Let $x_G$ be a random variable with mean $q$ and variance $\sigma_G^2 = q^2\delta$. In order to calculate and compare the moments of $x$ and $x_G$, it is useful to recall the moments of Gaussian- and Poisson-distributed variables:

$$\langle k^m \rangle = \sum_{j=0}^{m} q^j \left\{ {m \atop j} \right\} \tag{A.1}$$

$$\langle \psi^m \rangle = \sum_{j=0}^{\lfloor m/2 \rfloor} \frac{(-m)_{2j}\sigma^{2j}}{2^j j!} \tag{A.2}$$


*

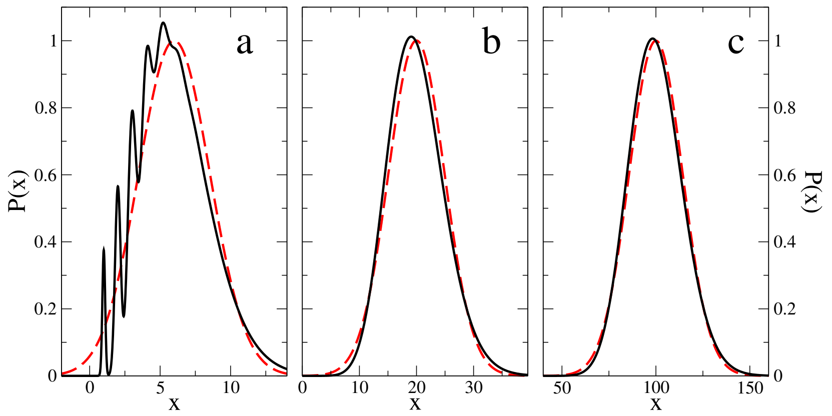


Figure 1: Comparison between $P(x)$ and its Gaussian approximation. Full curves give the true distribution $P(x)$ whereas dashed curves give the Gaussian approximation. Distributions are normalized to the peak of the corresponding Gaussian. a) $\delta = 0.1$, $q = 6$. b) $\delta = 0.1$, $q = 20$. c) $\delta = 0.1$, $q = 100$.

where $\left\{ {m \atop j} \right\}$ are Stirling numbers of the second kind, and $(-m)_j = (-m)(-m+1)\cdots(-m+j-1)$, $j > 0$ ($(-m)_0 = 1$). Using this, the moments of $x$ and $x_G$ are:

$$\begin{aligned} \frac{\langle x^m \rangle}{q^m} &= 1 + \frac{m(m-1)}{2}(\sigma^2 + 1/q) + \frac{m(m-1)}{4}\left(\frac{(m-2)(m-3)}{2}\sigma^4 + \right. \\ &\quad \left. + \frac{(m-2)(3m-5)}{6}q^{-2} + m(m-1)\sigma^2 q^{-1}\right) + \mathcal{O}(\delta^3) \qquad \text{(A.3)} \\ \frac{\langle x_G^m \rangle}{q^m} &= 1 + \frac{m(m-1)}{2}\delta + \frac{m(m-1)(m-2)(m-3)}{8}\delta^2 + \mathcal{O}(\delta^3) \qquad \text{(A.4)} \end{aligned}$$

Therefore, the difference between the moments is of order $\delta^2$ and both distributions converge as $\delta \to 0$. For small positive values of $\delta$, it can be assumed that the differences between averages calculated using these distributions will be of order $\delta^2$, if the functions are smooth enough [1]. For real systems, if photoreceptor noise is low, differences between the distribution become important when $q \lesssim 10$ (see Fig. 1).

### B Approximation for the Fisher information

For $n$ photoreceptor types, there are $n(n-1)/2$ possible channels that oppose all possible pairs of photoreceptors. The output of the $ij$ channel is modeled as:

$$f_{ij}^{opp} = \log f_i - \log f_j = \gamma \log O_{ij} \qquad \text{(B.1)}$$

with

$$O_{ij} = x_i / x_j \qquad \text{(B.2)}$$

It is important to note that, in general, the outputs of all the possible channels are not independent. For example, the output of opponent channel 13 can be obtained from 12 and 23 ($f_{13}^{opp} = f_{12}^{opp} - f_{23}^{opp}$). The largest set of independent channels can have at most $n-1$ channels.

As mentioned in the main text, the Fisher information does not change if a monotonous function of the responses. Thus, in order to simplify calculations, the variables $O_{ij}$ are used. Furthermore, for notational simplicity, it is better to use a rescaled version of the opponency channels output, defined as $\hat{O}_{SM} = O_{SM}q_M/q_S$ and $\hat{O}_{LM} = O_{SM}q_M/q_L$. Using the Gaussian approximation of the previous section, the probability distribution for $\hat{O}_{SM}$ and $\hat{O}_{LM}$ is

$$P(\hat{O}_{SM}, \hat{O}_{LM}) = \frac{h_1^2(\hat{O}_{SM}, \hat{O}_{LM}) + h_2(\hat{O}_{SM}, \hat{O}_{LM})}{2\pi\sqrt{\delta_S\delta_M\delta_L}\, h_2^{5/2}(\hat{O}_{SM}, \hat{O}_{LM})} \exp\left(-\frac{g(\hat{O}_{SM}, \hat{O}_{LM})}{h_2(\hat{O}_{SM}, \hat{O}_{LM})}\right) \tag{B.3}$$

where

$$\begin{aligned} g(\hat{O}_{SM}, \hat{O}_{LM}) &= \frac{(\hat{O}_{SM} - \hat{O}_{LM})^2}{\delta_S\delta_L} + \frac{(1-\hat{O}_{LM})^2}{\delta_M\delta_L} + \frac{(1-\hat{O}_{SM})^2}{\delta_M\delta_S} \\ h_k(\hat{O}_{SM}, \hat{O}_{LM}) &= \frac{1}{\delta_M} + \frac{\hat{O}_{LM}^k}{\delta_L} + \frac{\hat{O}_{SM}^k}{\delta_S} \quad (k=1,2), \end{aligned} \tag{B.4}$$

In the following it will be assumed that all the $\delta$ are of the same order. Given that the intrinsic noises of the photoreceptors are unlikely to differ by orders of magnitude, the only possibility for a $\delta$ to be much larger than the others could be for a wavelength for which $q_i(\lambda)$ is small. But in this case the corresponding photoreceptor is essentially inactive and $P(\hat{O}_{SM}, \hat{O}_{LM})$ reduces to the probability function for dichromats, and thus what follows still applies. Therefore, we will consider the expression "$\delta \to 0$" to mean that all $\delta$s tend to 0.

The inspection of Eq. B.5. shows that in the limit $\delta \to 0$ the function tends to 0 if $(\hat{O}_{SM}, \hat{O}_{LM}) \neq (1,1)$, and tends to infinity if $(\hat{O}_{SM}, \hat{O}_{LM}) = (1,1)$. Thus, $P(\hat{O}_{SM}, \hat{O}_{LM})$ becomes very concentrated around $(1,1)$ for small $\delta$ (see Fig. 2). This allows us to assume that $\hat{O}_{SM} = 1 + \delta^{\infty/\in}$ and $\hat{O}_{LM} = 1 + \delta^{\infty/\in}$.

$$\begin{aligned} \frac{\partial^2 \ln P(\mathbf{r}|\mathbf{s})}{\partial s^2} &= -\frac{g''}{2h_2} + \frac{gh_2''h_2/2 + (g'h_2 - gh_2')h_2'}{h_2^3} \\ &\quad -2\frac{q_M'}{q_M} + \frac{q_S'}{q_S} + \frac{q_L'}{q_L} - \frac{1}{2}(\frac{\delta_M'}{\delta_M} + \frac{\delta_L'}{\delta_L} + \frac{\delta_S'}{\delta_S}) - \frac{5}{2}\frac{h_2''h_2 - (h_2')^2}{h_2^2} \\ &\quad + \frac{(2(h_1')^2 + 2h_1h_1'' + h_2'')(h_1^2 + h_2) - (2h_1h_1' + h_2')^2}{(h_1^2 + h_2)^2} \end{aligned} \tag{B.5}$$

where ' indicates a derivative over $s$, and the dependencies of each function have not been included, for the sake of conciseness. In order to find the dominant term in the previous equation when $\delta$ is small, it is important to note that this implies that both $\sigma^2$ and $1/q$ are small. But in what follows we will not assume any special relationship between them. In other words we do not assume any particular value for $q\sigma^2$, nor for $q\delta$. We choose $s = I$, and use the definition of $q_i$ (Eq. 9 of the main text). The procedure for the case $s = \lambda$ is completely analogous. The calculation of the dominant part of each of the terms in Eq.B.5 is straightforward, but we give below a couple of examples.

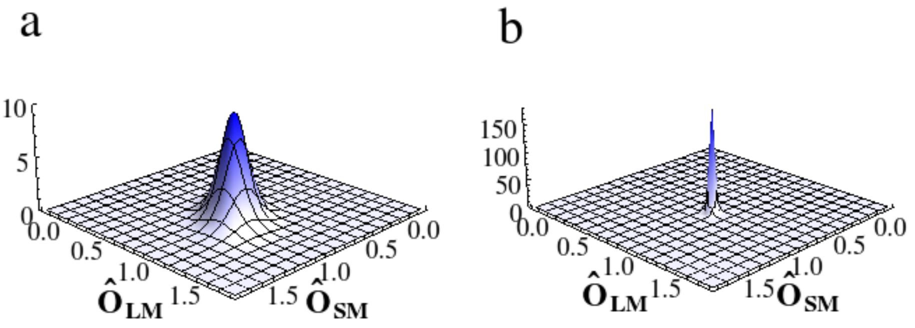


Figure 2: Probability distribution of rescaled opponency channel outputs $P(\hat{O}_{SM}, \hat{O}_{LM})$ for two different values of receptor noise. a) $\delta_S = \delta_M = \delta_L = 0.1$, b) $\delta_S = \delta_M = \delta_L = 0.02$.

Using the definition of $\delta$, we get $\delta' = -q'/q^2$. Since $q' = R(\lambda)$ which is a constant, we write $\delta' = \mathcal{O}(q^{-2})$ and, furthermore $\delta' = \mathcal{O}(q^{-3})$. Using the definition of $h_1$ in Eq.B.4, we get $h_1 = \mathcal{O}(\delta^{-1})$, $h_1' = \mathcal{O}((\delta q)^{-1})$, etc. In order to calculate the dominant term in $g$ it is important to remember that $\hat{O}_{SM}$ depends on $q$ (since it is the rescaled version of $O_{SM}$). Table 1 gives the dominant terms of all the functions involved in Eq.B.5.

| | $\delta'$ | $\delta''$ | $h_1$ | $h_1'$ | $h_1''$ | $h_2$ | $h_2'$ | $h_2''$ | $g$ | $g'$ | $g''$ |
|---|---|---|---|---|---|---|---|---|---|---|---|
| $\mathcal{O}$ | $\frac{1}{q^2}$ | $\frac{1}{q^3}$ | $\frac{1}{\delta}$ | $\frac{1}{q\delta}$ | $\frac{\sigma^2}{(q\delta)^2}$ | $\frac{1}{\delta}$ | $\frac{1}{q\delta}$ | $\frac{\sigma^2}{(q\delta)^2}$ | $\frac{1}{\delta}$ | $\frac{1}{\delta^{3/2} f}$ | $\frac{1}{(\delta f)^2}$ |

Table 1: Order of the dominant terms in each of the functions that compose the expression of the Fisher information.

Using this information in Eq.B.5, we find that the dominant term is the first summand, which is of order $1/(\delta f^2)$, whereas the dominant order of the rest is $1/f^2$. Thus, we get:

$$\left\langle \frac{\partial^2 \ln P(\mathbf{r}|\mathbf{s})}{\partial s^2} \right\rangle = -\frac{1}{2} \left\langle \frac{g''}{h_2} \right\rangle (1 + \mathcal{O}(\delta)). \tag{B.6}$$

If we now inspect the expression of $g''$ more closely, we see that it has several terms, three of which only one have order $1/(\delta f^2)$, whereas the most important term of the rest has order $1/(\delta^{1/2} f^2)$. Including this in the previous equation, gives

$$\left\langle \frac{\partial^2 \ln P(\mathbf{r}|\mathbf{s})}{\partial s^2} \right\rangle = \langle R(\hat{O}_{SM}, \hat{O}_{LM}) \rangle (1 + \mathcal{O}(\delta^{1/2})) \tag{B.7}$$

where

$$R(\hat{O}_{SM}, \hat{O}_{LM}) = \frac{\frac{1}{\delta_S \delta_L}(\hat{O}'_{LM}\frac{q'_L}{q_L} - \hat{O}_{SM}\frac{q'_S}{q_S})^2 + \frac{1}{\delta_S \delta_M}\hat{O}^2_{SM}(\frac{q'_L}{q_L} - \frac{q'_S}{q_S})^2 + \frac{1}{\delta_L \delta_S}\hat{O}^2_{LM}(\frac{q'_L}{q_L} - \frac{q'_M}{q_M})^2}{\delta_M^{-1} + \delta_S^{-1} O^2_{SM} + \delta_L^{-1} O^2_{LM}} \tag{B.8}$$

Now we prove that $\langle R(\hat{O}_{SM}, \hat{O}_{LM}) \rangle \to R(1,1)$ for vanishing $\delta$.

$$\begin{aligned} &|\langle R(\hat{O}_{SM}, \hat{O}_{LM}) \rangle - R(1,1)| \leq \langle |R(\hat{O}_{SM}, \hat{O}_{LM}) - R(1,1)| \rangle \\ &= \textstyle\int_{R^2} |R(\hat{O}_{SM}, \hat{O}_{LM}) - R(1,1)| P(\hat{O}_{SM}, \hat{O}_{LM}) dO_{SM} dO_{LM} \\ &= \textstyle\int_{\Omega} |R(\hat{O}_{SM}, \hat{O}_{LM}) - R(1,1)| P(\hat{O}_{SM}, \hat{O}_{LM}) dO_{SM} dO_{LM} \\ &+ \textstyle\int_{R^2-\Omega} |\langle R(\hat{O}_{SM}, \hat{O}_{LM}) - R(1,1)| P(\hat{O}_{SM}, \hat{O}_{LM}) dO_{SM} dO_{LM} \end{aligned} \tag{B.9}$$

where $\Omega = [1-\delta^*, 1+\delta^*] \times [1-\delta^*, 1+\delta^*]$. Using the Mean Value Theorem, we get, for the first integral:

$$\begin{aligned} &\int_{\Omega} |R(\hat{O}_{SM}, \hat{O}_{LM}) - R(1,1)| P(\hat{O}_{SM}, \hat{O}_{LM}) dO_{SM} dO_{LM} \\ &= \int_{\Omega} |\nabla R(\hat{O}^*_{SM}, \hat{O}^*_{LM})| |(\hat{O}_{SM}, \hat{O}_{LM}) - (1,1)| P(\hat{O}_{SM}, \hat{O}_{LM}) dO_{SM} dO_{LM} \\ &\leq \sqrt{2}\delta^* |\nabla R(\hat{O}^{max}_{SM}, \hat{O}^{max}_{LM})|. \end{aligned} \tag{B.10}$$

where we have used that $\nabla R$ reaches a maximum at some point $(\hat{O}^{max}_{SM}, \hat{O}^{max}_{LM}) \in \Omega$ because it is a continuous function. In order to bound the second integral, we need some bounds for $R$ and the exponential in the probability function $P(\hat{O}_{SM}, \hat{O}_{LM})$. Using some simple algebra, we obtain:

$$\begin{aligned} R(\hat{O}_{SM}, \hat{O}_{LM}) &\leq R^* = \frac{\max(R^*_S, R^*_L)}{\min(\delta_S^{-1}, \delta_L^{-1})} \\ R^*_i &= \frac{1}{\delta_S \delta_L}\left(\frac{(q'_S)^2}{q_S^2} + 2\frac{q'_S q'_L}{q_S q_L}\right) + \frac{1}{\delta_S \delta_M}\left(\frac{q'_S}{q_S} - \frac{q'_M}{q_M}\right)^2 \end{aligned} \tag{B.11}$$

for $i = S, L$, and

$$\frac{g(\hat{O}_{SM}, \hat{O}_{LM})}{h_2(\hat{O}_{SM}, \hat{O}_{LM})} \geq C^* \frac{\delta^{*2}}{\delta_M} = \frac{\min(\delta_S^{-1}, \delta_L^{-1})}{8\max(\delta_S^{-1}, \delta_L^{-1}) + \delta_M^{-1}} \frac{\delta^{*2}}{\delta_M} \tag{B.12}$$

for $(\hat{O}_{SM}, \hat{O}_{LM}) \in R^2 - \Omega$. Thus, we obtain

$$\begin{aligned} & \int_{R^2-\Omega} |\langle R(\hat{O}_{SM}, \hat{O}_{LM}) - R(1,1)| P(\hat{O}_{SM}, \hat{O}_{LM}) dO_{SM} dO_{LM} \\ \leq \;& \exp(-C^* \frac{\delta^*}{\delta_M})(R^* + R(1,1)) \int \frac{h_1^2(\hat{O}_{SM}, \hat{O}_{LM}) + h_2(\hat{O}_{SM}, \hat{O}_{LM})}{2\pi\sqrt{\delta_S \delta_M \delta_L}\, h_2^{5/2}(\hat{O}_{SM}, \hat{O}_{LM})} dO_{SM} dO_{LM} \\ \leq \;& \exp(-C^* \frac{\delta^{*2}}{\delta_M})(R^* + R(1,1))(3 + \delta_S^{-1} + \delta_L^{-1} + \delta_M^{-1}))/3 \end{aligned} \tag{B.13}$$

The calculations above are valid for arbitrary $\delta^* > 0$. Consider now the case $\delta^* = \delta^{1/2-\epsilon}$, for any positive $\epsilon$. Excluding the exponential, the terms in the bound of the above equation form a rational function of $\delta$. Thus, in the limit of vanishing $\delta$ the bound above tends to 0 faster than any polynomial in $\delta$. Now, it is easy to see that the bound found for the first integral (Eq.B.10) is a polynomial in $\delta$, and therefore it becomes the dominant term in the limit of vanishing $\delta$. Furthermore, since $|\nabla R(\hat{O}_{SM}^{max}, \hat{O}_{LM}^{max})|$ is of the same order as $R(1,1)$ (i.e. $\mathcal{O}(\delta^{-1}q{-}2)$), we finally obtain

$$|\langle R(\hat{O}_{SM}, \hat{O}_{LM})\rangle - R(1,1)| \leq C\delta^{1/2-\epsilon} R(1,1) \tag{B.14}$$

The bound obtained is probably rather loose, as shown by the fact that already for realistic values of $\delta$, $R(\hat{O}_{SM}, \hat{O}_{LM}))\rangle$ and $R(1,1)$ are very close (see Fig.1 in the main text).

### C Approximation of a rescaled product of Gaussian variables

Since in all this section we are dealing with photoreceptors of the same class, for the sake of conciseness in the following the subindex $i$ will be dropped. Let $x_1$,...,$x_n$ be random variables drawn from the same Gaussian distribution with mean $q$ and variance $q^2\delta^2$. We are interested in the probability distribution of the random variable

$$x^* = (x_1 * \cdots * x_n)^{1/a}. \tag{C.1}$$

Noticed that this is completely equivalent to calculating the probability distribution of

$$x^* = q^{n/a}(x_1 * \cdots * x_n)^{1/a}, \tag{C.2}$$

where the $x_i$ are now drawn from a Gaussian with mean 1 and variance $\delta^2$. Thus in the following we approximate a product of $n$ such random variables. To go back to $x^*$ one needs only to rescale it with $q^{n/a}$. For a Gaussian random variable we have [2, 3]:

$$\langle |x|^a \rangle = \frac{2^{a/2}}{\Gamma(\frac{a+1}{2})}\, {}_1F_1\left(-\frac{a}{2}; \frac{1}{2}; -\frac{1}{2\delta}\right), \tag{C.3}$$

where ${}_1F_1(a;b;z)$ is Kummer's confluent hypergeometric function. Using for this function an expansion for large values of $|z|$ [4], we get:

$$\langle |x|^a \rangle = \sum_{k=0}^{k^*} \frac{(-a)_{2k}\sigma^{2k}}{2^k k!}, \tag{C.4}$$

where $k^*$ is the optimal truncation index, which is usually large ($k^* \sim 1/\delta$) [4]. In Eq. C.4 we have discarded terms that are exponentially small (or, more precisely, terms of type $\mathcal{O}(\exp(a/\delta)p(\delta))$, where $p(\delta)$ is any polynomial in $\delta$. This allows us to develop the moments of $x$ in powers of $\delta$:

$$\langle (x^*)^m \rangle = 1 - \frac{mn}{2}a(1-ma)\delta + \frac{mn}{8}a(1-ma)(man-6+(4-man)ma)\delta^2 + \mathcal{O}(\delta^3). \tag{C.5}$$

Let $x^*_G$ be a random variable drawn from a Gaussian distribution with mean $\langle x^*_G \rangle = \langle x^* \rangle$ and variance $\langle x^*_G \rangle^2 \delta_G$, with

$$\delta_G = a^2 n\delta[1 + (5 + a(an-6))\frac{\delta}{2} + \mathcal{O}(\delta^2)]. \tag{C.6}$$

Using Eq. A.2, the moments of $x^*_G$ can be readily calculated. It can then be checked that

$$\langle (x^*)^m \rangle - \langle (x^*_G)^m \rangle = \frac{a^3 n}{3} m(m-1)(m-2)(an-1)\delta^2. \tag{C.7}$$

This shows that the distributions of $x^*$ and $x^*_G$ converge. Note that, according to the equation above, for the particular case of $a = 1/n$ the differences between moments should be even smaller. And, indeed, a similar (but lengthier) calculation as above for $a = 1/n$ gives:

$$\langle (x^*)^m \rangle - \langle (x^*_G)^m \rangle = \frac{7(n-1)}{n^3} m(m-1)(m-2)\delta^3. \tag{C.8}$$

This implies that in the limit of vanishing $\delta$ the Gaussian approximation is much better in the limit of vanishing $\delta$. Fig 3 shows that this seems also to be the case of realistic values of $\delta$.

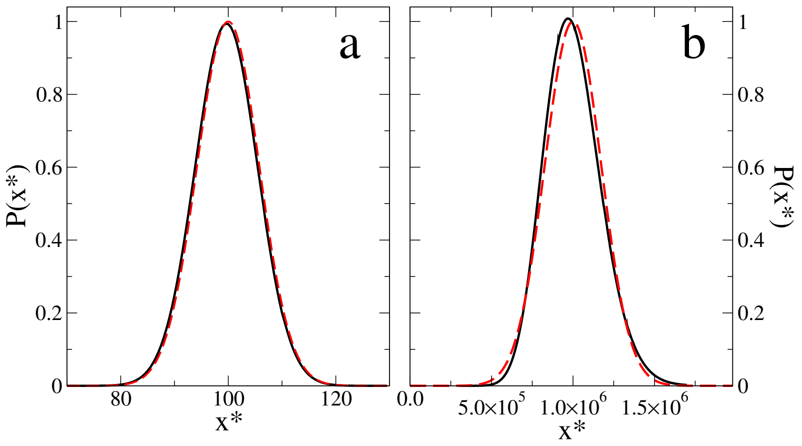


Figure 3: Comparison between $P(x^*)$ and its Gaussian approximation. Full curves give the true distribution whereas dashed curves give the approximations. Distributions are normalized to the peak of the corresponding Gaussian. a) Purely chromatic, $n = 3$, $q = 100$, $\delta = 0.01$. b) Non purely chromatic, $a = 1$, $n = 3$, $q = 100$, $\delta = 0.01$ .
